# Efficiency of *Salicornia neei* to treat aquaculture effluent from a hypersaline and artificial wetland

**DOI:** 10.1101/2020.09.06.259358

**Authors:** Mónica R. Diaz, Javier Araneda, Andrea Osses, Jaime Orellana, José A. Gallardo

## Abstract

In this study we evaluated the potential of *Salicornia neei*, a halophyte plant native to South America, to treat saline effluents with simulated concentration of ammonium-N (Amm) and nitrate-N (Nit) similar to land-based marine aquaculture effluents. Plants were cultivated for 74 days in drainage lysimeters under three treatments of seawater fertilized with: 1) Nit+Amm, 2) Nit, or 3) without fertilizer (Control). Over 5 repetitions, nitrogen removal efficiency (RE) was high in both treatments (Nit + Amm = 89.6± 1,0 %; Nit 88.8 ± 0.9 %). While nitrogen removal rate (RR) was non linear and concentration-dependent (RR_day 1-4_: Nit+Amm= 2.9 ± 0.3 mg L^−1^ d^−1^, Nit = 2.4 ± 0.5mg L^−1^ d^−1^; RR_day5-8_: Nit + Amm = 0.8 ± 0.2mg L^−1^ d^−1^, Nit=1.0 ± 0.2mg L^−1^ d^−1^). Effluent salinity increased from 40.6 to 49.4 g L^−1^ during the experiment, with no observed detrimental effects on RE or RR. High nitrogen removal efficiency and significant biomass production observed, Nit+Amm = 11.3 ± 2.0 kg m^−2^; Nit = 10.0 ± 0.8 kg m^−2^; Control = 4.6 ± 0.6 kg m^−2^, demonstrate that artificial wetlands of *S. neei* can be used for wastewater treatment in saline aquaculture in South America.

## 1. Introduction

Aquaculture provides nearly 50% of the world’s fish production, and it is expected to increase to 60% by 2030 due to the growing demand for marine fishery products [1]. Land-based marine aquaculture systems will play an important role in meeting this demand and will also do so in a more environmentally sustainable way regarding marine aquaculture in the ocean [2, 3]. However, the development of marine recirculating aquaculture systems (RAS) is limited by the ability to efficiently treat saline wastewater, which accumulates a large amount of nitrogen compounds derived from the metabolism of culture organisms [3–5]. In these RAS, the removal of nitrogen compounds, mainly ammonium (NH_4_^+^) and ammonia (NH_3_^−^), becomes a priority for elimination because they quickly deteriorate the water quality and cause negative effects on the culture [6, 7]. Biofilters that promote the conversion of ionized and deionized ammonium to nitrate (NO_3_^−^) are usually used for this purpose [8, 9]. NO_3_^−^is not very toxic to most cultured organisms [10, 11], with tolerable accumulated concentrations reported between 120 mg L^−1^ of NO_3_^−^ and 150 mg L^−1^ of NO_3_^−^ in marine RASs [12].

Recent developments of integrated systems allow the use of RAS waste products as nutrients, coupling different water loops with the main fish production water system [13]. To take advantage of these waste products, such as nitrogen compounds that accumulate in marine RAS, the use of artificial wetlands with facultative or obligate halophytes has been proposed [14–16]. Halophyte plants have the ability to absorb different forms of N, depending on different environmental factors such as the availability of CO_2_ [17]. For example, some species of the genus *Spartina* show a higher affinity for NH_4_^+^ consumption [18, 19], while others like *Juncus maritimus*, have a marked preference for NO_3_^−^, even in substrates that contained high availability of NH_4_^+^ [20]. Also, if the plants are grown in lysimeters or wetland, the interaction with soil, microorganism and plant have a higher potential to remove nitrogen compounds and produce biomass, which can be used as animal feed or human food [21, 22], and in the production of biofuels or by-products of interest to the pharmaceutical industry [2, 5, 15, 23, 24], among others. Additionally, it has been demonstrated that these systems are also efficient in removing residual phosphates from RASs [2, 15, 23, 25–27].

*Salicornia neei* is a succulent hydrohalophyte of herbaceous habit, native to South America and abundantly distributed on the South Pacific coast, where much of the marine aquaculture production in South America is concentrated [28]. *S. neei* is used as a gourmet food and is a type of emerging crop in the coastal zone of Chile. This plant has been described as containing high amounts of nutrients and important functional metabolites [22]. Additionally, physiological studies have been performed to observe germination patterns [29] and changes in the concentration of metabolites and antioxidants when exposed to different salinity gradients [30].

The objective of this study was to evaluate the capacity of the halophyte *S. neei* for use as a sink for dissolved nitrogen compounds in effluent from land-based marine aquaculture systems and to simultaneously evaluate the resulting biomass production. The data obtained in this study will allow us to establish whether *S. neei* is a plant suitable for treating land-based marine aquaculture effluent with the potential for use in marine recirculating aquaculture systems.

## 2. Materials and Methods

### 2.1. Collection of plant material and acclimatization

In July 2014, 100 *Salicornia neei* plants with fully developed roots and shoots were collected in the “Salinas de Puyalli” wetland, located in the commune of Papudo, Valparaíso Region, Chile (32° 24′ 54″ S, 71° 22′ 43″ W) and subsequently transferred to the “Laboratorio Experimental de Acuicultura” of the Pontificia Universidad Católica de Valparaíso, in Valparaíso, Chile (33° 1′ 21″ S, 71° 37′ 57″ W). Plants were sown in sand beds and irrigated with Hoagland solution once a week for 10 weeks. Once the plants adapted and recovered their vigour, they were transferred to the experimental unit.

### 2.2. Experimental unit

The experimental unit consisted of three RAS separated to each other, each one composed of three drainage lysimeters (replicates). Each lysimeter was housed in a polyethylene container measuring 0.5 m × 0.6 m × 0.6 m (length × width × depth) with a surface area of 0.3 m^2^ and a total area per RAS of 0.9 m^2^. In each lysimeter, four *S. neei* plants were planted until reaching a biomass of approximately 1 kg per lysimeter or 3 kg m^−2^. A leachate collection system was installed in each lysimeter, consisting of a perforated pipe at the bottom to collect the water, followed by a layer of gravel with a diameter of 0.5 cm and height of 15 cm and polyethylene mesh with 0.3 mm pore size to cover the gravel. For the substrate, coarse sand was used until reaching 35 cm high (Figure 1). Each RAS was connected to a nutrient storage, which in turn was fed by a main tank that contained filtered seawater. Each nutrient storage tank was equipped with an aeration pump to promote biological nitrification processes. The irrigation water supply (influent) was performed with a 0.5 HP centrifugal pump (Humboldt, TPM60). Each RAS was supplied daily with 30 litres m^−2^ d^−1^ of water through a drip irrigation system, programmed to run for 15 minutes at 09:00 and at 17:00 hrs. This guaranteed that a large proportion of the irrigation water will penetrate and be collected to the bottom of the lysimeter. Drainage water (effluent) was returned to the respective collection tanks of each system to close the recirculating water loop.

**Figure 1.**
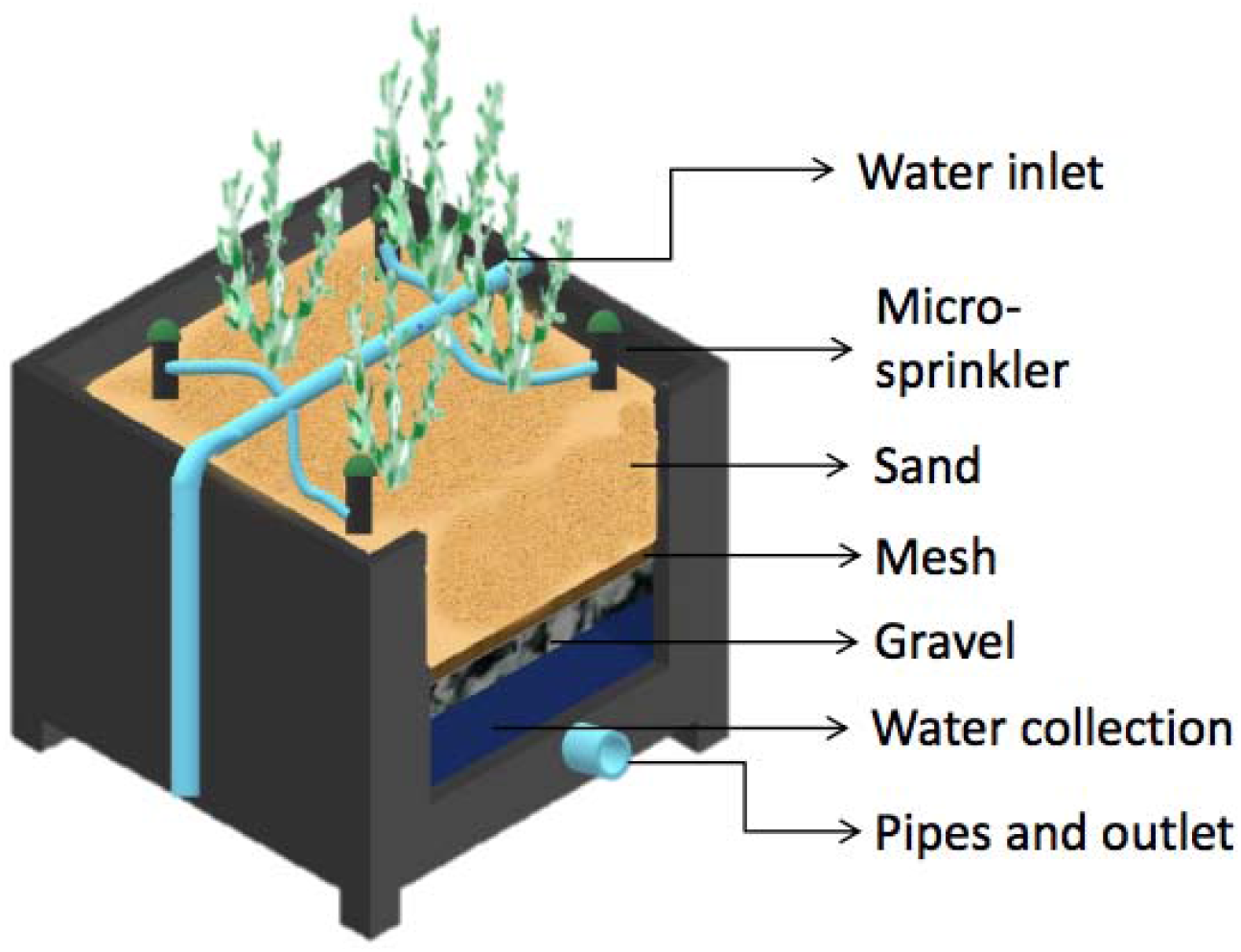
The diagram shows the design of one lysimeter, depicting the overall construction, water inlet and outlet, substrate (sand and gravel separated by a mesh), and irrigation micro-sprinklers.

### 2.3. Experimental design

The *S. neei* performance regarding removal of nitrogen compounds and biomass production was evaluated for 74 days under three irrigation treatments: 1) seawater fertilized with nitrate-N + ammonium-N (Nit+Amm); 2) seawater fertilized with nitrate-N (Nit); and 3) seawater without fertilizer that was used as a control group (Control). The nutrient concentrations in each irrigation water supply were designed according to the typical average concentrations of ammonium-N (NH_4_^+^-N) and nitrate (NO_3_^◻^N) reported in land-based marine aquaculture effluent [31, 32]. The following concentrations were used: Nit+Amm = 1 mg L^−1^ of TAN (total ammonia nitrogen) and 100 mg L^−1^ of NO_3_^◻^N; Nit = 100 mg L^−1^ of NO_3_^◻^N; and Control = no fertilizer. The nutrient solution for each RAS was prepared directly in each collection tank and was completely renewed every 14-15 days. Five repetitions or Inputs were performed during the 74 days of culture.

The physico-chemical parameters of water quality were recorded directly from the drainage water during the first eight consecutive days after nutrient addition. The estimation of NO_3_^◻^N concentration was performed using the cadmium reduction method. NO_3_^◻^N removal efficiency (RE) was calculated as: RE = (Ci − Co)/Ci ⍰ 100 where: Ci = concentration of NO_3_^◻^N in the influent water at day 1; Co = concentration of NO_3_^◻^N in the effluent water at day 8 from each input. Additionally, temperature, oxygen, conductivity, salinity and pH were measured as water quality indicators. These parameters were measured using a HACH multiparameter probe (HQ40). Biomass (fresh weight) was recorded at the beginning and at the end of the experiment using a scale (Jadever, JWE-6K). The data on ambient temperature, rainfall and relative humidity were sourced from climate records of the Chilean Meteorological Office (Torquemada-Viña del Mar Station).

### 2.4. Statistical analysis

Biomass was compared using a two-way ANOVA in R Statistical Software [33], with interaction between nitrogen simulated concentrations (3 levels: Nit + Amm, Nit and Control) and days of culture (two levels: 0 and 74 days)(Supplementary table S1). To compare groups means we performed post hoc Tukey tests (Supplementary table S2) in R Statistical Software [33]. The change in Nitrate-nitrogen (NO_3_^L^N) concentration showed a negative non linear relationship through the crop, therefore the linear removal rate (RR) was calculated separately for days 1 to 4 and days 5 to 8 of each Input, when linearility was observed. Linear regression models with respond (dependent) variable Nitrate-nitrogen (NO_3_^L^N) concentration and predictor variables days of culture and nitrogen simulated concentrations were performed using the “lm” function in R Statistical Software [33]. Probabilities of p<0.05 were considered significant.

## 3. Results

### 3.1. Environmental conditions and RAS parameters

During the 74 days of culture, the ambient temperature and relative humidity conditions and the temperature, pH and salinity of the cultivation system showed different levels of variability, and no rainfall was recorded during the experiment. The ambient temperature had a mean of 16 ± 4 °C but was highly variable during the day with extreme values of 9 and 31 °C, while the relative humidity was 77.8 ± 8.7%, with extreme values of 60% and 95% (Supplementary figure S1). The temperature in the culture systems was usually higher than the ambient temperature, with a mean of 20.5 ± 1.24 °C and a range of 19.1 to 21.7 °C, with no observed differences between treatments (Table 1). The pH remained relatively constant and without differences between treatments, while the salinity had a noticeable increase from a mean of 40 g L^−1^ of NaCl on day 1 to a mean of 51.5 ± 0.19 g L^−1^ of NaCl at the end of the experiment (Table 1). No significant differences in salinity between treatments were observed (p<0.05).

**Table 1.** RAS physicochemical parameters measured on the effluent. Temperature, pH, and salinity (mean ± SE) recorded at the effluent of the culture systems with *Salicornia neei*. Salinity is expressed as gram of natrium chloride per liter (g L^−1^). Treatments were irrigated with nitrate-N and ammonium-N (Nit + Amm), nitrate-N (Nit) or with sea water only (Control group). Mean values of 3 lysimeters per treatment are displayed (± EE).

| Input | Treatment | Temperature<br>($^{\circ}\text{C}$) | pH | Salinity<br>($\text{g L}^{-1}$) |
| --- | --- | --- | --- | --- |
| 1 | Nit + Amm | $18.2 \pm 4.2$ | $8.2 \pm 0.1$ | $40.6 \pm 2.2$ |
| | Nit | $19.5 \pm 4.7$ | $8.2 \pm 0.1$ | $41.3 \pm 1.9$ |
| | Control | $19.1 \pm 4.3$ | $8.2 \pm 0.1$ | $40.0 \pm 0.0$ |
| 2 | Nit + Amm | $18.8 \pm 1.6$ | $8.1 \pm 0.1$ | $44.9 \pm 2.3$ |
| | Nit | $21.7 \pm 3.3$ | $8.1 \pm 0.1$ | $48.4 \pm 2.2$ |
| | Control | $18.6 \pm 1.5$ | $8.0 \pm 0.1$ | $43.6 \pm 2.1$ |
| 3 | Nit + Amm | $20.8 \pm 0.6$ | $7.9 \pm 0.1$ | $48.5 \pm 2.5$ |
| | Nit | $21.2 \pm 0.8$ | $7.9 \pm 0.1$ | $48.8 \pm 3.2$ |
| | Control | $20.8 \pm 0.5$ | $8.0 \pm 0.1$ | $43.6 \pm 2.1$ |
| 4 | Nit + Amm | $20.2 \pm 1.2$ | $8.0 \pm 0.1$ | $47.5 \pm 1.9$ |
| | Nit | $20.6 \pm 1.4$ | $8.0 \pm 0.1$ | $47.5 \pm 2.1$ |
| | Control | $20.3 \pm 1.2$ | $8.2 \pm 0.1$ | $46.5 \pm 2.6$ |
| 5 | Nit + Amm | $20.6 \pm 0.6$ | $8.0 \pm 0.1$ | $48.0 \pm 2.2$ |
| | Nit | $20.9 \pm 0.7$ | $7.9 \pm 0.1$ | $47.7 \pm 2.4$ |
| | Control | $20.7 \pm 0.5$ | $8.2 \pm 0.1$ | $46.5 \pm 1.6$ |

### 3.2. Growth and biomass formation

Regarding biomass production, the treatments with Nit+Amm and Nit showed a significant increase in fresh weight from 3.0 ± 0.6 g to 11.3 ± 2.0 kg m^−2^ and from 3.4 ± 0.1 g to 10.0 ± 0.8 kg m^−2^, respectively (Figure 2). Although the plants grew in the control group, this increase in biomass was not significant (P= 0.61).” Plants irrigated with seawater presented chlorosis and accumulation of pigment in leaf tissue, probably anthocyanin, which indicated moderate stress on the plant, however, this phenomenon was not observed in either of the two nitrogen treatments (Figure 3).

**Figure 2.**
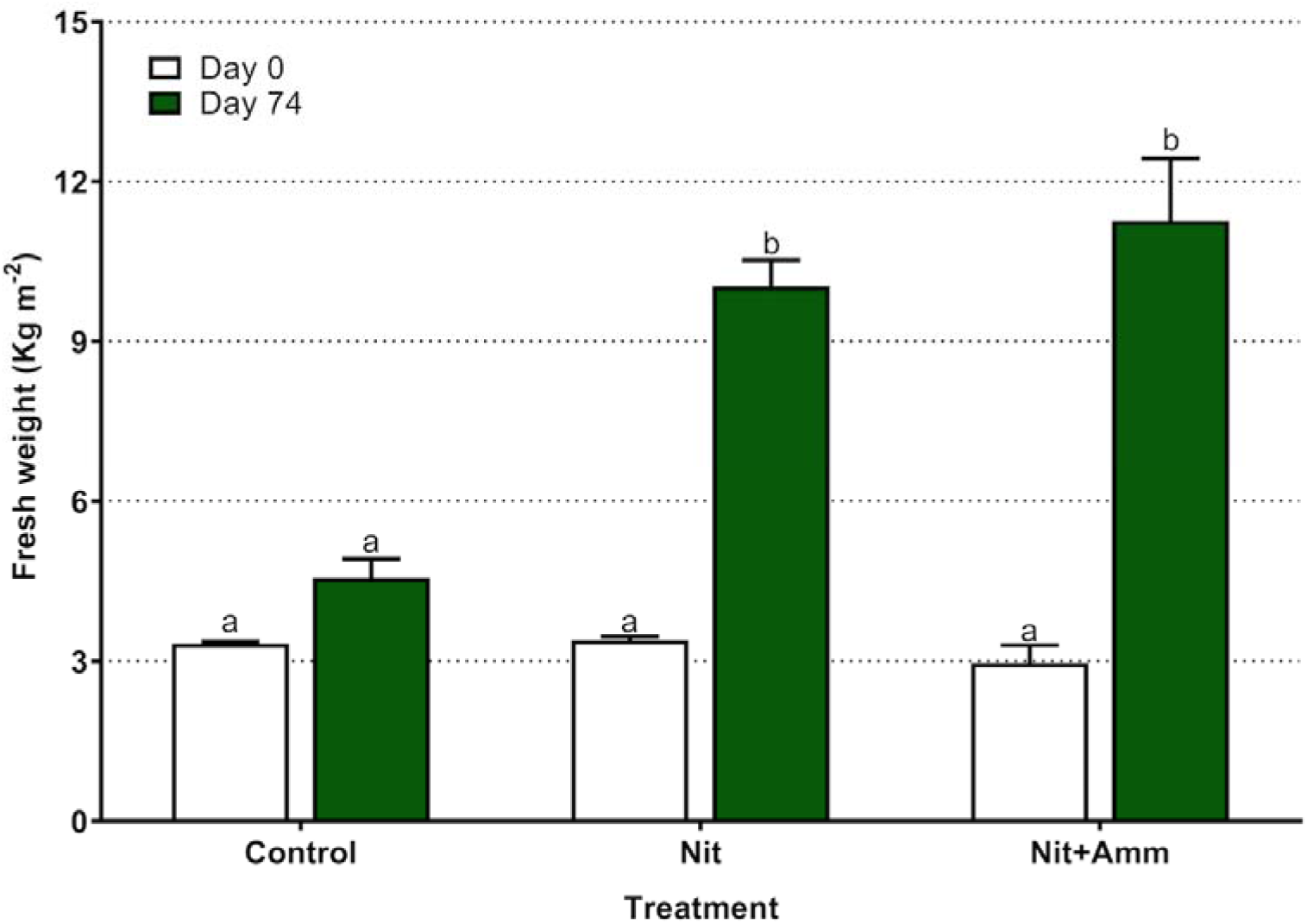
Production of biomass of *Salicornia neei* by treatment expressed as yield of fresh weight per area unit (kg m^−2^). Nit + Amm: corresponds to the treatments irrigated with nitrate-N and ammonium-N, Nit: irrigated with nitrate-N, Control: treatment irrigated with sea water only. Lower-case letters represents significant differences between treatments. Mean values of 3 lysimeters per treatment are displayed (± EE).

**Figure 3.**
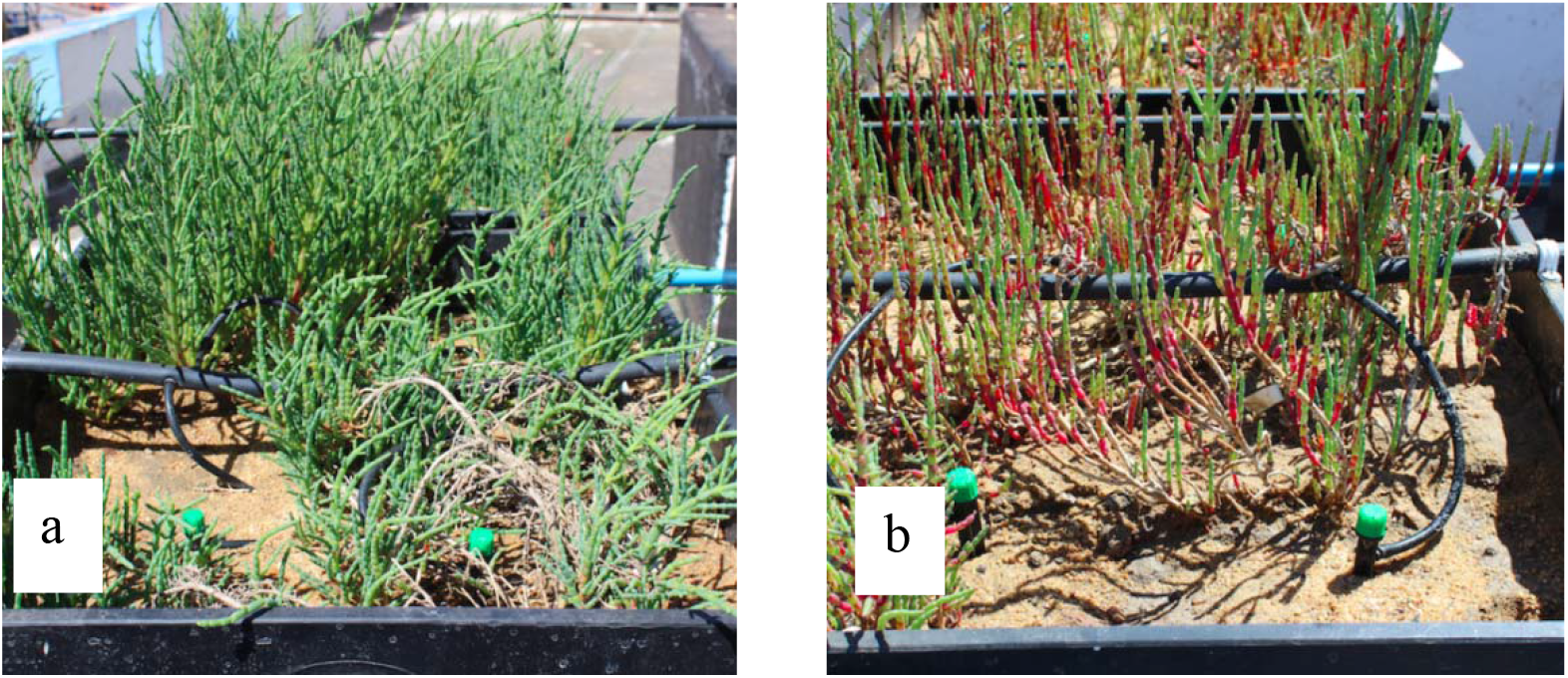
Picture of two lysimeters with Salicornia neei at the end of the experiment (day 74). Picture (a) plants irrigated with nitrate and ammonium (Nit + Amm). Picture (b) plants irrigated with sea-water (control).

### 3.3 Efficiency of Salicornia neei to treat saline effluent

Nitrate-N removal was non linear and concentration-dependent for treatments Nit + Ammand Nit (Figure 4). Thus, Nitrate-N removal rates (RR) were reducing when reducing nitrogen loading from 2.9 ± 0.3 mg L^−1^ d^−1^ (RR_day 1-4_) to 0.8 ± 0.2mg L^−1^ d^−1^ (RR_day5-8_) in the treatment Nit +Amm, and from 2.4 ± 0.5 mg L^−1^ d^−1^ (RR_day 1-4_) to 1.0 ± 0.2mg L^−1^ d^−1^ (RR_day5-8_) in the treatment Nit (Table 2). On the other hand, Nitrate-N removal rates measured between days 1 to 4 (RR_day 1-4_) had a clear tendency to increase as biomass production increased at the treatment Nit but not in the treatment Nit + Amm (Table 2), which is perhaps explained by the greater availability of nitrogen in this last treatment. Without considering these differences in both treatments, the nitrogen removal efficiency was high in each treatment, in and throughout the crop, and varied between 87% and 92% (Table 2). Effluent salinity increased from 40.6 to 49.4 g L^−1^ during the experiment, with no observed detrimental effects on the Nitrate-N removal rates or on the nitrogen removal efficiency.

**Figure 4.**
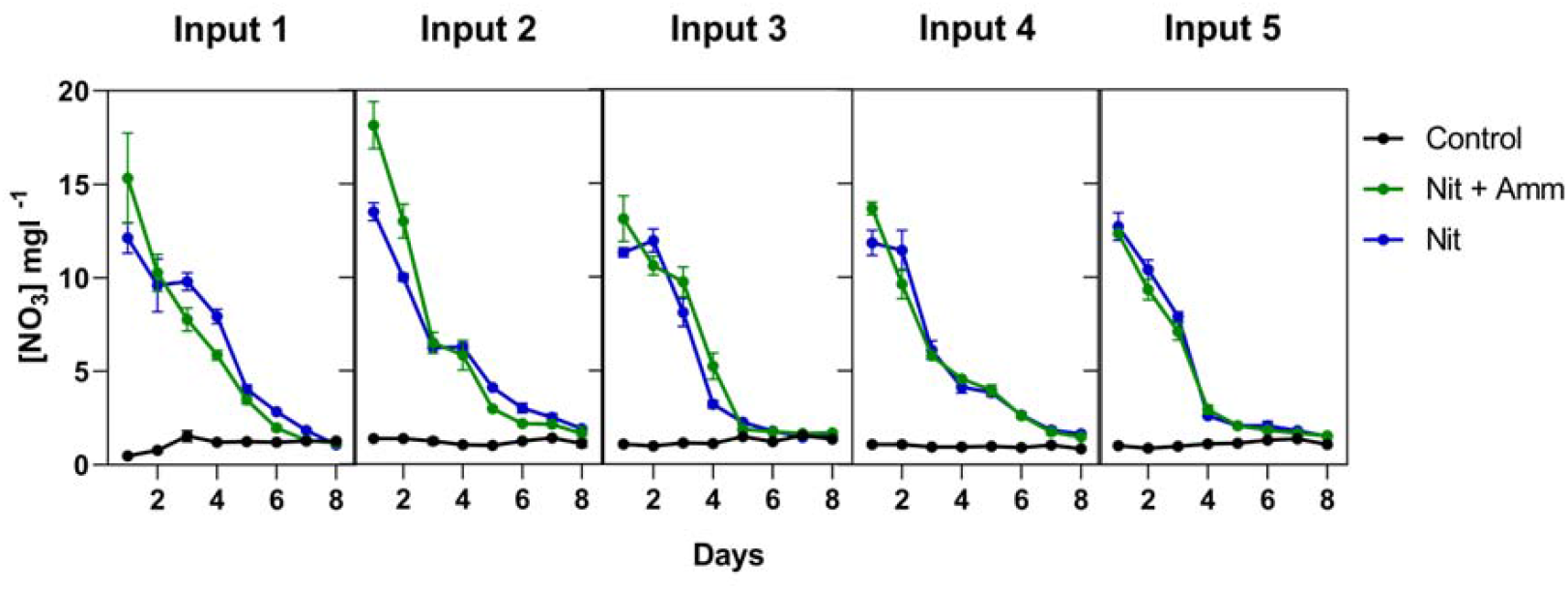
Nitrogen removal by treatment and input. Nitrate-nitrogen (NO_3_^◻^N) concentration in each treatment was expressed in mg L^−1^ and measured during 8 days from nutrient input over 74 days of experimentations. Treatments were irrigated with nitrate-N and ammonium-N (Nit + Amm), nitrate-N (Nit) or with sea water only (Control group). Mean values of 3 lysimeters per treatment are displayed (± SE).

**Table 2.** Nitrate-nitrogen (NO_3_ N) concentration in the influent water at day 1 (Ci) and in the effluent at day 8 (Co), removal efficiency (RE) and removal rate (RR) for each treatment irrigated with nitrate-N and ammonium-N (Nit + Amm) and nitrate-N (Nit). Each Input has 3 lysimeters per treatment. Treatments were irrigated with nitrate-N and ammonium-N (Nit + Amm) and nitrate-N (Nit). Mean values are displayed (± SE).

| Treatment | Input | Ci<br>($\text{mg L}^{-1}$) | Co<br>($\text{mg L}^{-1}$) | RE<br>(%) | RR<br>day 1-4<br>($\text{mg L}^{-1} \text{d}^{-1}$) | RR<br>day 5-8<br>($\text{mg L}^{-1} \text{d}^{-1}$) |
| --- | --- | --- | --- | --- | --- | --- |
| Nit + Amm | 1 | $14.0 \pm 4.2$ | $1.2 \pm 0.1$ | 91.4 | $2.9 \pm 0.5$ | $1.2 \pm 0.1$ |
| | 2 | $17.7 \pm 2.2$ | $1.8 \pm 0.7$ | 89.8 | $2.0 \pm 0.3$ | $0.3 \pm 0.1$ |
| | 3 | $13.5 \pm 0.6$ | $1.1 \pm 0.05$ | 91.9 | $3.1 \pm 0.3$ | $0.8 \pm 0.1$ |
| | 4 | $12.9 \pm 2.1$ | $1.7 \pm 0.1$ | 86.8 | $3.6 \pm 0.5$ | $0.0 \pm 0.0$ |
| | 5 | $12.4 \pm 0.1$ | $1.5 \pm 0.1$ | 87.9 | $3.0 \pm 0.2$ | $0.0 \pm 0.0$ |
| | Mean | | | $89.6 \pm 1.0$ | $2.3 \pm 0.4$ | $0.5 \pm 0.2$ |
| Nit | 1 | $12.0 \pm 1.4$ | $1.1 \pm 0.1$ | 90.8 | $1.1 \pm 0.5$ | $1.4 \pm 0.1$ |
| | 2 | $13.3 \pm 0.8$ | $1.8 \pm 0.2$ | 86.5 | $1.2 \pm 0.3$ | $0.6 \pm 0.1$ |
| | 3 | $12.4 \pm 1.2$ | $1.2 \pm 0.15$ | 90.3 | $2.8 \pm 0.3$ | $0.7 \pm 0.1$ |
| | 4 | $13.2 \pm 1.0$ | $1.6 \pm 0.1$ | 87.9 | $3.7 \pm 0.5$ | $0.2 \pm 0.0$ |
| | 5 | $13.0 \pm 1.3$ | $1.5 \pm 0.0$ | 88.5 | $3.3 \pm 0.2$ | $0.0 \pm 0.0$ |
| | Mean | | | $88.8 \pm 0.9$ | $2.9 \pm 0.5$ | $0.4 \pm 0.2$ |

## 4. Discussion

The integration of halophytes as a biofilter in recirculating systems in marine aquaculture has been proposed as an adequate alternative to decontaminating waters with increased nitrogen compounds [34]. In this study we evaluated if artificial wetlands of *S. neei* could be used to treat saline aquaculture effluent. *S. neei* was selected mainly due to its natural occurrence throughout much of the South Pacific coast of South America [35], which would allow its rapid adoption in the growing South American aquaculture. Nitrate-nitrogen removal rate and removal efficiency recorded in this study (Table 2) was higher or similar than those reported with other halophyte species in high salinity [14, 15]. Thus, artificial wetlands of *S. neei* could a good alternative to the treatment of highly concentrated wastewater released in marine RAS.

Physicochemical parameters of the effluent, such as temperature and pH are especially important in the treatment of saline wastewater because they can affect the determinant processes in the removal of nitrogen compounds [36]. In this study, temperature and pH were maintained within the optimal ranges (20-21 °C and 7.8-8.2) and therefore did not affect the nutrient removal processes (Table 1). This finding is consistent with Lee et al. [37], who reported that, for denitrification processes in wetland systems, the optimal temperature ranges between 20 and 40 °C and the optimal pH is approximately 8.0. Another important parameter evaluated in this study was the high effluent salinity, which reached concentrations of up to 50 g L^−1^ of NaCl. This increase was mainly due to the known environmental factor of evapotranspiration, consistent with a study by Freedman et al. [38], who found increased salinity of treated water in artificial wetlands despite the salt uptake by plants due to soil evaporation and plant transpiration.

Nitrogen bioaccumulation was not determined empirically in this study, but we derived it from Riquelme et al. [22], a previous study performed by our research group. Riquelme et al. [22] show that the total of N fixed in the aerial part of *S. neei* corresponds to 1.76 ± 0.08 g per 100 g of fresh weight. Similar results were obtained in *S. brachiata* by Rathore et al. [39] from India. Thus, we estimated that the total concentration of nitrogenous nutrients fixed in *S. neei* at the end of the trial would be between 46 and 103.9 g for the Nit treatment. While for Amm + Nit, the oscillatory fixation between 57.8 and 130.1 g of N for the total biomass formed by this treatment, indicating that *S. neei* could assimilated most of the nitrogen available in this test. According to these results, it can also be suggested that *S. neei* could store ammonium −N, if the differences of the estimate in the two treatments are considered (approximately 20% more N with the Amm + Nit treatment). This being a reflection of the synergy produced by these two compounds when consumed at the same time [40]. However some researchers currently believe that the actual absorption may represent only a relatively small fraction of the global rate of nitrogen (N) elimination [41] and microorganisms that play the most important role in the use and transformation of nitrogen component [42].

In response to this uncertainty, other researchers have studied and obtained low removal rates by plants. Specifically, Tanner et al. [43] found that of the total nitrogen removed by planted wetland systems, only 25% corresponded to fixation in plants. Likewise, Lin et al. [32] observed that of the 73% of nitrogen removed, only 11% had been fixed in plants. Notwithstanding the above, Webb et al. [25] observed significant differences between the nitrogen removal capacity in beds planted with and without halophytes. In their study, they demonstrated a higher removal yield in planted beds (62.0 ± 34.6 mmol N m^−2^ d^−1^) than in unplanted beds (23.0 ± 26.8 mmol N m^−2^ d^−1^). And this is consistent, with our results that show a nitrogen removal proportional to biomass. Therefore, we cannot rule out that the increase in biomass exclusively explains the increase in the nitrate removal rate. In fact, it is plausible that a strong root system formed by this class of plants supports the establishment of certain microorganisms that improve the removal rate of nitrogen loads by acting synergistically.

The formation of *S. neei* biomass during the evaluation period reached a total net weight of 7 - 8 kg m^−2^ over a period of eleven weeks in the treatments irrigated Nit and Nit+Amm respectively. These high yields in biomass production are comparable to those obtained by Ventura et al. [44], whose yields for *Salicornia persica* reached 16 kg m^−2^ in a span of 24 weeks. On the other hand, *S. neei* plants remained vigorous throughout the evolution period, even at high salinity concentrations close to 50 g L^−1^ of NaCl. This inherent feature of halophytes highlights the powerful response mechanisms to abiotic stress triggered by *S. neei*, reinforcing the feasibility of including this plant for aquaculture effluent treatment. Regarding removal of the two sources of nitrogen compounds, there was a positive interaction between the ammonium/nitrate supplied for biomass formation of *S. neei.* This positive interaction could be caused by the contribution of the nitrate ion that would act as an important osmotic anion for expansion of the foliar cells [45].

## 5. Conclusions

Our results reveal that the integration of *S. neei* into artificial wetlands with recirculating aquaculture effluent would be a viable alternative for eliminating nutrient loads in saline wastewater and that this plant could be included in marine RASs in South America.

## Author Contributions

All co-authors have fully participated in and accept responsibility for the work. This publication is approved by all authors and the responsible authorities where the work was carried out. The authors declare that they have no competing interests, and ensure that the work was appropriately investigated, resolved, and documented in the literature.

Conceptualization, J.A.G. and J.O.; methodology, J.A.G. and J.O.; software, M.R.D.; validation, J.A.G. and M.R.D.; formal analysis, M.R.D.; investigation, M.R.D., A.O., J.A.; data curation, M.R.D; writing—original draft preparation, J.A.G. and M.R.D..; writing—review and editing, J.A.G. and J.O.; visualization, M.R.D.; project administration, J.A.G.; funding acquisition, J.A.G. and J.O.. All authors have read and agreed to the published version of the manuscript

## Acknowledgments

The authors wish to thank Mr. Aldo Madrid and Marine Farms Inc. for his technical assistance and support during the project. M.D. was supported by a Doctoral fellowship by the “Dirección General de Investigación y Postgrado” of the Universidad Técnica Federico Santa María.

