## Supplementary material for "Efficiency of *Salicornia neei* to treat aquaculture effluent from a hypersaline and artificial wetland": Suplemental Material

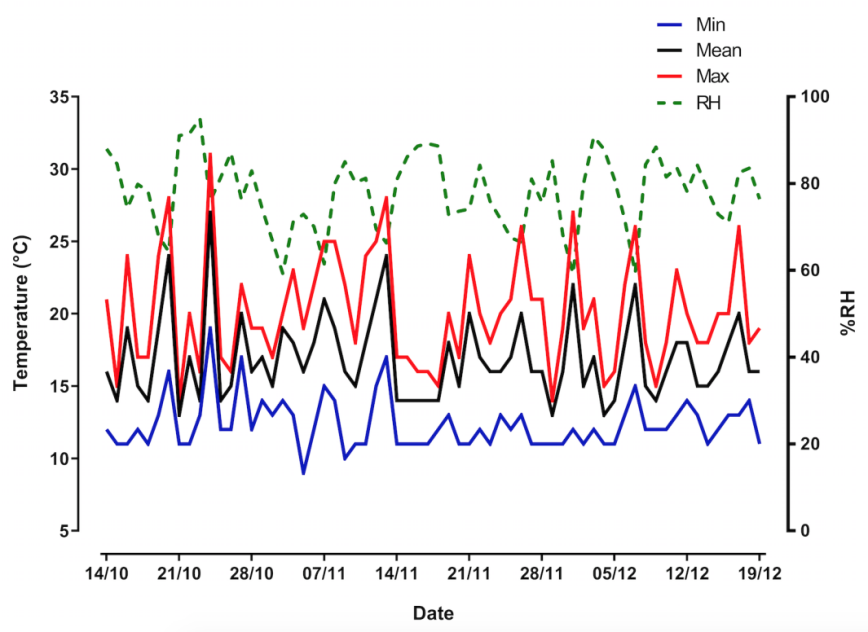

**Figure S1.** Ambient temperature (°C) and relative humidity (%RH) during the date of experimentation. The graphic shows mean, maximum and minimum values for the ambient temperature, over 74 days.

**Table S1.** Two-way ANOVA. For quantitative biomass growth data, the effect of Treatment versus Days_of_culture were analyzed by two-way ANOVA test.

| Two-way ANOVA | Degree of freedom | Sum of Squares | Mean of Squares | F Value | Probability |
| --- | --- | --- | --- | --- | --- |
| Treatment | 2 | 35.58 | 17.79 | 19.55 | 0.000168 |
| Days_of_culture | 1 | 131.13 | 131.13 | 144.14 | 4.81e-08 |
| Treatment:Days_of_culture | 2 | 40.86 | 20.43 | 22.46 | 8.78e-05 |
| Residuals | 12 | 10.92 | 0.91 |  |  |

**Table S2.** Tukey test (HSD). Tukey multiple comparisons of means 95% family-wise confidence level.

|  | Mean differences | lower | uper | p adj |
| --- | --- | --- | --- | --- |
| Con:0-Amm + Nit:0 | 0.36722222 | -2.248671 | 2.983115 | 0.9963521 |
| Nit:0-Amm + Nit:0 | 0.43144444 | -2.184449 | 3.047338 | 0.9923336 |
| Amm + Nit:74-Amm + Nit:0 | 8.30844444 | 5.692551 | 10.924338 | 0.0000021 |
| Con:74-Amm + Nit:0 | 1.61411111 | -1.001782 | 4.230004 | 0.3608018 |
| Nit:74-Amm + Nit:0 | 7.07077778 | 4.454885 | 9.686671 | 0.0000117 |
| Nit:0-Con:0 | 0.06422222 | -2.551671 | 2.680115 | 0.9999993 |
| Amm + Nit:74-Con:0 | 7.94122222 | 5.325329 | 10.557115 | 0.0000034 |
| Con:74-Con:0 | 1.24688889 | -1.369004 | 3.862782 | 0.6128292 |
| Nit:74-Con:0 | 6.70355556 | 4.087662 | 9.319449 | 0.0000204 |
| Amm + Nit:74-Nit:0 | 7.87700000 | 5.261107 | 10.492893 | 0.0000037 |
| Con:74-Nit:0 | 1.18266667 | -1.433226 | 3.798560 | 0.6601556 |
| Nit:74-Nit:0 | 6.63933333 | 4.023440 | 9.255226 | 0.0000226 |
| Con:74-Amm + Nit:74 | -6.69433333 | -9.310226 | -4.078440 | 0.0000207 |
| Nit:74-Amm + Nit:74 | -1.23766667 | -3.853560 | 1.378226 | 0.6196403 |
| Nit:74-Con:74 | 5.45666667 | 2.840774 | 8.072560 | 0.0001604 |
